# Phosphatase-mediated mitigation of rare earth element toxicity to *Pseudomonas putida*

**DOI:** 10.64898/2026.08.03.742403

**Authors:** Charly A. Dupont, Théophile Franzino, Lola Perrey, Maximilien Beuret, Nolhan Berceaux, Patrick Billard

## Abstract

Anthropogenic activities are driving an increasing flux of rare earth elements (REE) into environmental compartments, raising concerns about their biological impact, particularly on microorganisms that sustain ecosystem functioning. Here, we provide a systematic assessment of the toxicity of all 16 REE toward *Pseudomonas putida* KT2440, a soil bacterium that can use these metals as enzyme cofactors. Dose-response growth inhibition assays revealed high sensitivity to light REE. Toxicity correlated strongly with ionic radius, with IC_50_ values ranging from 0.3 µM for lanthanum to 10 µM for scandium. Serial propagation of *P. putida* under gradually increasing REE stress yielded resistant populations, from which two stably resistant strains were isolated. Genome resequencing showed that both strains carried a single mutation in *uxpB*, encoding an alkaline phosphatase. Gene deletion and overexpression experiments, together with phosphatase activity measurements, confirmed the involvement of *uxpB* in REE resistance. Our findings reveal a previously unrecognized mechanism of tolerance to REE, suggesting that mutations enhancing phosphatase activity promote phosphate release from organic phosphorus compounds and REE immobilization, thereby mitigating toxicity.

## 1. Introduction

Rare earth elements (REE) are a group of metals with similar chemical properties comprising the series of 15 lanthanides (Ln; Z=57 to 71), together with scandium (Z=21) and yttrium (Z=39). Based on their atomic mass and geochemical behavior, they are commonly divided in two categories, light (LREE, La to Eu) and heavy (HREE, Gd to Lu). Their chemical similarities are largely due to their common trivalent oxidation state (+3), the small differences in atomic number, and comparable ionic radius [1]. A key factor influencing these properties is the lanthanide contraction, a progressive decrease in ionic radii with increasing atomic number caused by the stronger nuclear attraction of the outer electrons [2]. This phenomenon results in subtle but significant variations in their chemical behavior and industrial applications. Indeed, the unique electronic configuration of REE endows them exceptional electronic, magnetic, and optical properties, making them indispensable in modern technologies, from permanent magnets for wind turbines and electric motors, to catalysts and medical imaging. Despite their name, REE are not truly rare in the Earth’s crust and widely occur together in phosphate, carbonate, or silicate minerals. Their natural concentration are comparable to those of biologically important micronutrients such as Zn, Cu or Ni [3,4]. The LREE have significantly higher crustal abundances than HREE, with average concentrations varying from 66 mg.kg^-1^ (Ce) to 0.4 mg.kg^-1^ (Lu). In soils, the cumulative concentrations of all REE typically range between 100 and 1,000 mg.kg^-1^ [5].

Due to their wide applications in high-tech products, demand for these metals, and therefore their extraction and use are steadily increasing [6]. This has inevitably led to rising levels of REE in both soils and waters in areas close to large cities or industrial complexes [5,7,8], as well as in agricultural areas, especially in China where they have long been applied as REE-enriched phosphate fertilizers [5,9]. Over the last decades, growing concerns about emerging REE pollution and its implications for ecosystem and human health has stimulated an increasing number of studies investigating their fate, bioavailability, and ecotoxicological effects across diverse ecosystems. A large body of literature has reported adverse effects of REEs for diverse groups of organisms [8,10–12], as well as REE-associated hormetic effects, i.e. stimulatory responses at low doses and inhibitory responses at high doses, notably in plants [13]. In comparison, far fewer studies have focused on toxicity to bacteria, even though they are recognized as drivers of key ecosystem processes and can act as early sentinels of environmental perturbations. Furthermore, data on the element-specific toxicity of REEs and on the molecular mechanisms underlying bacterial responses to these metals are still scarce [11,14,15].

In contrast, the toxicological effects of other metals (most notably cadmium, zinc, and copper) toxicity have been extensively characterized. At elevated concentrations, some metals can catalyze the formation of reactive oxygen species and impair the electron transport chain. They can also compete with essential trace metals, thereby reducing their cellular uptake. In addition metals may inactivate metalloenzymes and regulatory proteins by replacing the native metal ion in their active sites through mismetallation [16]. To avoid such outcomes, bacteria have evolved a variety of mechanisms to ensure metal homeostasis. A first system involves small molecules, including organic and inorganic ligands or specific metal-binding proteins that buffer the cytoplasmic free metal concentration at relevant range despite high overall concentration [16–18]. If metal concentration exceeds the buffering capacity, dedicated efflux systems can be mobilized to pump the excess free ions outside the cells [19]. Bacteria are also capable of immobilizing metals through extracellular sequestration or periplasmic retention, for example by releasing ligands such as phosphate or sulfide that precipitate metals into less toxic forms [20,21].

Due to their insolubility, REEs have long been regarded as non-essential for living organisms, and their biological significance rarely considered. This assumption was overturned in the 2010s by the discovery of REE-dependent enzymes in methylotrophic [22,23] and non methylotrophic bacteria [24], leading to the emergence of the fast-moving field of REE biology [25,26]. Despite advances in understanding the mechanisms by which these bacteria acquire REE, little attention has been paid to their possible toxic effects when metabolized. It is worth noting that the culture media commonly used are highly complexing, promoting the formation of insoluble species notably with phosphate, carbonates and hydroxides, meaning that their toxicity could be underestimated [29]. Therefore, chemical speciation modelling using freeware geochemical tools such as Visual MINTEQ or PHREEQC is essential for estimating free ion concentrations in growth media and predicting toxicity.

The soil-dwelling bacterium *Pseudomonas putida* KT2440 employs LREE as a cofactor for a periplasmic pyrroloquinoline quinone (PQQ)-dependent alcohol dehydrogenase (ADH) and appears to possess a complete machinery dedicated to the sensing, uptake, and utilization of these metals [24,27,28]. Here we investigate how *P. putida* manages exposure to toxic concentrations of REE under growth conditions where these metals are not used for its metabolism. Dose-response growth inhibition assays were first performed in a defined growth medium optimized to maximize REE bioavailability and to permit speciation modelling. These analyses provided a comprehensive assessment of toxicity patterns across a set of 16 REE and enabled the identification of potential relationships between their toxic effects and physicochemical properties. An adaptive laboratory evolution experiment was then performed to better understand the mechanisms underlying tolerance to REE. Serial propagation of *P. putida* under gradually increasing REE stress levels yielded resistant populations from which two strains showing a stable and significantly higher resistance were isolated. Sequencing of their genomes revealed that each evolved strain has a single substitution in the same *uxpB* gene encoding and alkaline phosphatase. Gene deletion and overexpression experiments together with phosphatase activity measurements confirmed involvement of *uxpB* in REE resistance. Furthermore, we report how single mutations in evolved isolates enhance tolerance through phosphatase-mediated REE immobilization.

## 2. Results

### 2.1. Toxicity of rare earth elements toward *Pseudomonas putida* KT2440

Dose-response growth inhibition assays were performed to determine the susceptibility of *P. putida* to the 16 REE by calculating IC_50_ values after 24 hours of growth in a low metal-complexing medium. The results revealed that LREE (La to Eu) were more toxic than HREE (Gd to Lu, excluding Sc and Y) under these conditions, as indicated by their IC_50_ values ranging from 0.31 µM to 0.69 µM, compared to IC_50_ values between 0.96 µM and 1.59 µM for the latter (Table 1 and Figure S1). This observation is supported by a Pearson correlation coefficient of 0.93 between ionic radii and IC_50_ values, showing a strong relationship between those parameters. By contrast, Sc and Y did not follow this trend, with IC_50_ values of 10.05 and 0.61 µM, respectively, reducing the correlation between ionic radii and IC_50_ to 0.77. Since time-dependent decrease in REE exposure concentrations in the test medium can occur [29], we compared the effective concentrations of representative REE in the MOPS medium immediately after amendment and after 24 hours of incubation. The measured concentrations proved to be comparable to the nominal values and remained unchanged after incubation, regardless of the metal (Figure S2). Thus, the varying toxicity of individual REE to *P. putida* is unlikely to result from abiotic alterations in their bioavailability. For comparison purposes, growth inhibition tests were also performed with Cd, Zn and Ni. The highest toxicity was observed for Cd, followed by Ni and Zn (Table 2 and Figure S3). The IC_50_ values calculated for these three metals were much higher, by about two to three orders of magnitude, than those for REE, confirming the high sensitivity of *P. putida* to the latter.

**Table 1:** IC_50_ values of 16 REE for *P. putida* KT2440. Growth inhibition tests were conducted in MOPS medium with glycerol 1-phosphate as the phosphorus source. The concentrations of REE tested ranged from 0 to 10 µM. A line separates light and heavy REE. Data were obtained after 24h of incubation at 28°C and 180 rpm in biological triplicate with technical duplicate (n=3). IC_50_ values were calculated using a four-parameter log-logistic non-linear regression based on the dose response curves presented in supplementary Figure S1. ^†^: Ionic radii obtained from (Shannon, 1976). ^††^: ^61^Pm was not tested due to its radioactivity.

| **Ionic radius (Å)<sup>†</sup>** | **Element<sup>††</sup>** | **IC<sub>50</sub> (μM)** | **95% CI (μM)** |
| --- | --- | --- | --- |
| 1.032 | <sup>57</sup>La | 0.31 | 0.28 to 0.35 |
| 1.01 | <sup>58</sup>Ce | 0.38 | 0.30 to 0.51 |
| 0.99 | <sup>59</sup>Pr | 0.65 | 0.55 to 0.77 |
| 0.983 | <sup>60</sup>Nd | 0.36 | 0.27 to 0.50 |
| 0.958 | <sup>62</sup>Sm | 0.68 | 0.62 to 0.75 |
| 0.947 | <sup>63</sup>Eu | 0.69 | 0.63 to 0.76 |
| 0.938 | <sup>64</sup>Gd | 1.01 | 0.85 to 1.18 |
| 0.923 | <sup>65</sup>Tb | 1.08 | 0.87 to 1.37 |
| 0.912 | <sup>66</sup>Dy | 0.96 | 0.79 to 1.16 |
| 0.901 | <sup>67</sup>Ho | 1.29 | 1.13 to 1.51 |
| 0.89 | <sup>68</sup>Er | 1.27 | 1.08 to 1.68 |
| 0.88 | <sup>69</sup>Tm | 1.16 | 1.06 to 1.29 |
| 0.868 | <sup>70</sup>Yb | 1.12 | 0.90 to 1.34 |
| 0.861 | <sup>71</sup>Lu | 1.59 | 1.23 to 2.31 |
| 0.745 | <sup>21</sup>Sc | 10.05 | 9.85 to 10.32 |
| 0.90 | <sup>39</sup>Y | 0.61 | 0.60 to 0.62 |

| Element | WT* |  | EvoNd5 |  | p-value |
| --- | --- | --- | --- | --- | --- |
|  | IC<sub>50</sub> (μM) | 95% CI (μM) | IC<sub>50</sub> (μM) | 95% CI (μM) |  |
| <sup>30</sup>Zn | 1.19 | 1.08 to 1.31 | 2.00 | 1.74 to 2.32 | <0.0001 |
| <sup>28</sup>Ni | 0.89 | 0.83 to 0.96 | 1.29 | 1.17 to 1.45 | 0.0604 |
| <sup>48</sup>Cd | 0.28 | 0.26 to 0.29 | 0.33 | 0.29 to 0.38 | 0.0017 |

### 2.2. Identification of adaptive mutations conferring improved tolerance to REE

To explore the adaptive response of *P. putida* KT2440 to REE, serial transfers of cultures with increasing concentrations of NdCl_3_ from 25 nM to 2 µM were performed (Figure 1A), followed by single colony isolation from endpoint tolerant populations. Two evolved strains, designated EvoNd5 and EvoNd6, exhibited stable tolerance to high metal concentration with IC_50_ values increased by 10.7-and 7.7-fold respectively compared to the WT* strain (Figure 1B, Figure S4, Table S1). Whole genome sequencing of these two strains showed single point mutations in the *uxpB* gene (PP_1043; Figure 1C) encoding an alkaline phosphatase of the PhoX family [30]. The mutations led to G5S and G47S substitutions, both located in the peptide leader of UxpB, for the EvoNd5 and EvoNd6 strains, respectively (Figure 1C). The emergence of two independent mutations in the same region of UxpB suggests the enzyme may act as a key mediator of the adaptive response to REE.

**Fig. 1.**
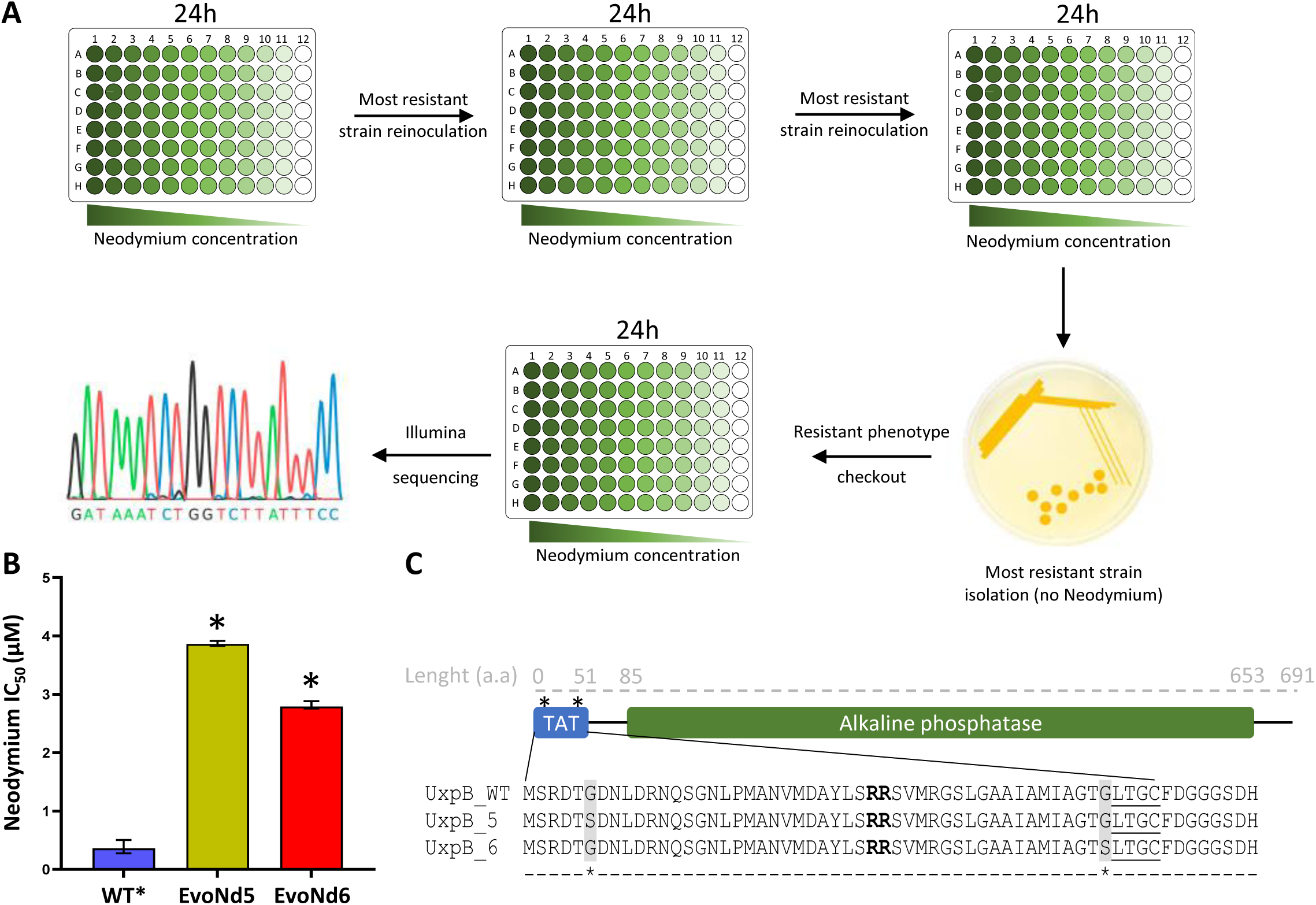
Adaptive laboratory evolution identified the alkaline phosphatase encoding gene *uxpB* as a putative player in REE stress-resistance. (A) Schematic overview of the adaptive laboratory evolution experiment performed with increasing Nd concentrations. Resistant endpoint populations were isolated and screened for stable Nd tolerance before genome sequencing. (B) Nd IC_50_ values of the evolved strains EvoNd5 and EvoNd6 compared with the parental WT* strain. **** p < 0.0001. (C) Mutations identified in evolved strains. Both strains carried a single amino acid substitution in UxpB (PP_1043), highlighted in grey. Predicted Tat signal peptide and alkaline phosphatase domains are indicated. The twin arginine and lipobox motifs are indicated in bold and underlined, respectively.

### 2.3. Contribution of the UxpB alkaline phosphatase to REE stress tolerance

Deletion of *uxpB* abolished growth on MOPS medium supplemented with glycerol 1-phosphate (Figure 2A), indicating that this phosphatase is required for utilizing this phosphorus source. This however prevented assessment of the effects of the *uxpB* gene deletion on REE toxicity in our conditions. To overcome this limitation, the deletion strain was transformed with the pJN105 plasmid carrying the wild-type *uxpB* gene (Δ*uxpB*+*uxpB* strain) under the control of the L-arabinose inducible *araBAD* promoter. Even in the absence of L-arabinose, this construct partially restored the growth of the Δ*uxpB* strain (Figure 2A). Under these conditions, both the lag phase and the maximal OD_600_ (OD_600_max) at stationary phase were higher than those of the control strain carrying the empty pJN105 plasmid (WT*+EV). When the medium was supplemented with 0.0125% L-arabinose, which slightly induces the *araBAD* promoter [31], the lag phase of the Δ*uxpB*+*uxpB* strain became comparable to that of the control strain, while its growth rate and its OD_600_max were slightly higher. Upon strong induction with 0.1% L-arabinose, the lag phase was further reduced, and both growth rate and OD_600_max were similar to those observed under the 0.0125% L-arabinose condition (Figure 2A). L-arabinose had no detectable effect on the growth of the control strain (data not shown).

**Fig. 2.**
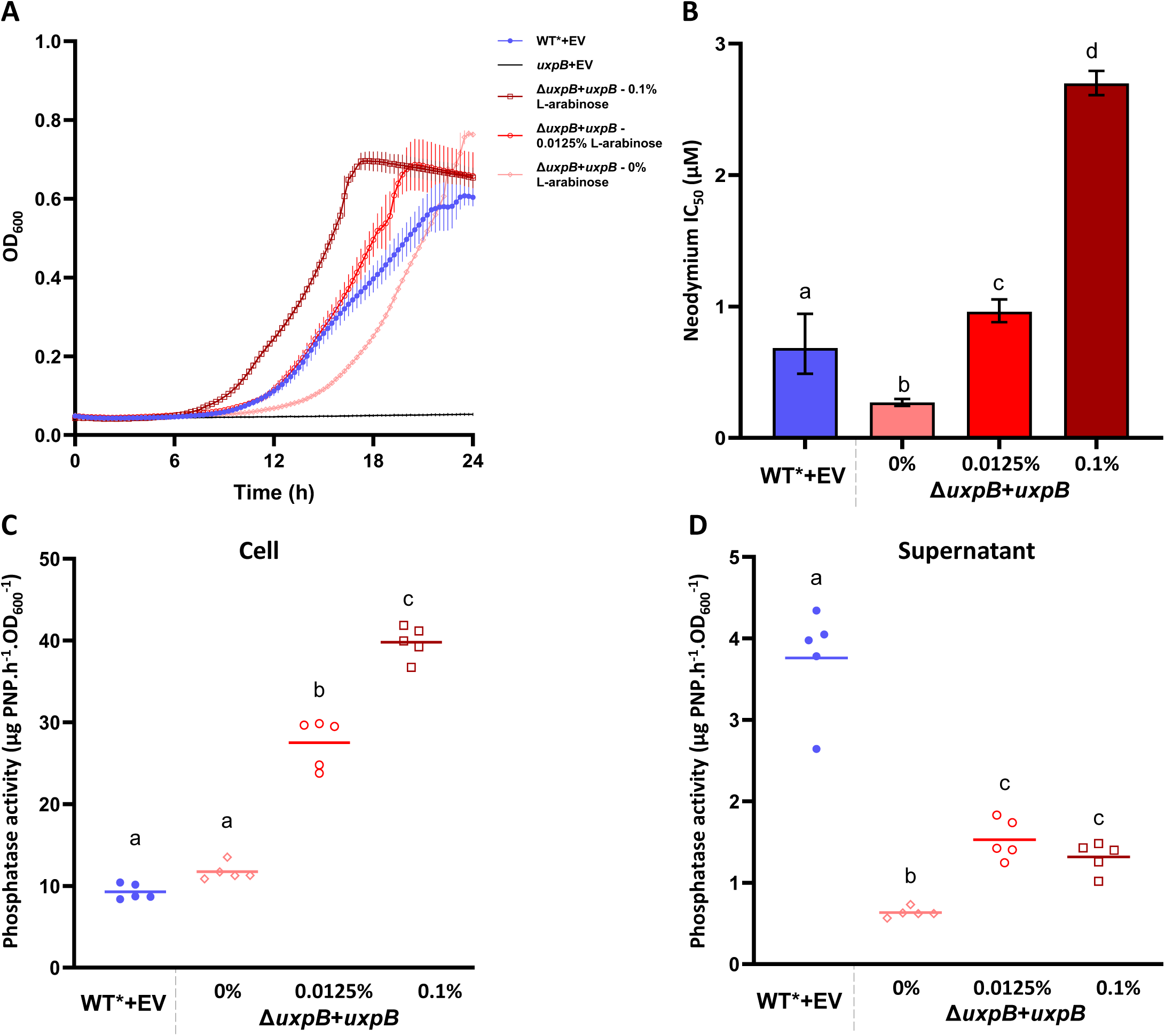
UxpB is involved in REE resistance. (A) Growth of WT* and Δ*uxpB* strains carrying either the empty pJN105 vector (EV) or a pJN105 plasmid expressing *uxpB* (Δ*uxpB*+*uxpB*) in MOPS medium containing glycerol 1-phosphate. *uxpB* expression was induced with 0, 0.0125, or 0.1% (m/v) L-arabinose. Data represent means ± SD from three biological replicates with technical duplicates (n = 3). (B) Nd IC_50_ values of WT*+EV and complemented Δ*uxpB*+*uxpB* strains under different induction conditions. Error bars represent 95% confidence intervals. Different letters indicate significant differences (extra-sum-of-squares F test, p < 0.05; n = 3). (C-D) Cell-associated (C) and extracellular (D) phosphatase activities of WT*+EV and Δ*uxpB*+*uxpB* strains after 24 h of growth. Activity was measured by pNPP hydrolysis after 1h and normalized to OD_600_. Points represent biological replicates and bars indicate means (n = 5). Different letters indicate significant differences (one-way ANOVA with Tukey’s post hoc test, p < 0.05).

The toxicity of Nd to the different strains was then evaluated by determining their IC_50_ values under the same conditions. The IC_50_ of Nd for the control strain (WT*+EV) was 0.68 µM. In the absence of *uxpB* induction, the Δ*uxpB*+*uxpB* strain was more sensitive to Nd, showing a 2.5-fold decrease in IC_50_ (0.27 µM) compared to the control. When *uxpB* expression was induced, Nd tolerance increased, with IC_50_ values 1.4-fold higher in the presence of 0.0125% L-arabinose (0.96 µM) and 4-fold higher with 1% L-arabinose (2.70 µM) relative to the control strain (Figure 2B, Figure S5, Table S1). These results indicate that UxpB expression confers enhanced tolerance to Nd in a dose-dependent manner.

Alkaline phosphatase activity assays indicated that the control WT*+EV cells produced approximately 10 µg of p-nitrophenol (PNP) per hour and per OD_600_ unit from a 4 mM p-nitrophenyl phosphate solution. In the corresponding supernatant, an activity of about 3.8 µg PNP·h^-^¹·OD ^-^¹ was detected (Figure 2C, 2D). For the Δ*uxpB*+*uxpB* strain without L-arabinose induction, the cell-associated activity was comparable to that of the control strain, while the supernatant activity dropped to 0.7 µg PNP·h^-^¹·OD_600_^-^¹ - a 5.4-fold decrease (Figure 2C, 2D). Upon addition of 0.0125% L-arabinose, cell-associated phosphatase activity increased 2.8-fold relative to the control (28 µg PNP·h^-^¹·OD ^-^¹), whereas the supernatant activity reached 1.5 µg PNP·h^-^¹·OD ^-^¹, 2.5 times lower than that of the control strain (Figure 2C, 2D). When 0.1% L-arabinose was added, cell-associated activity rose further to 38 µg PNP·h^-^¹·OD ^-^¹ (3.8-fold increase), while supernatant activity remained similar to that observed under 0.0125% induction (Figure 2C, 2D). From these data we conclude that UxpB is mainly cell-associated and that its expression level correlates with total phosphatase activity

### 2.4. Single-point mutations in UxpB alter its localization and activity

Cell-associated phosphatase activity in EvoNd5 and EvoNd6 increased by 3.9- and 3-fold, respectively, compared to the WT* strain (Figure 3A). In the supernatant, alkaline phosphatase activity was approximately eight times higher in both evolved strains than in the WT* strain (Figure 3B). When grown with glycerol 1-phosphate as the sole phosphorus source, the evolved strains exhibited a 3-hour reduction in lag phase, while growth rates and OD_600_max remained almost unchanged (Figure 3C). It therefore appears that point mutations in the leader peptide of UxpB improve both the cell-associated and the supernatant-associated alkaline phosphatase activity.

**Fig. 3.**
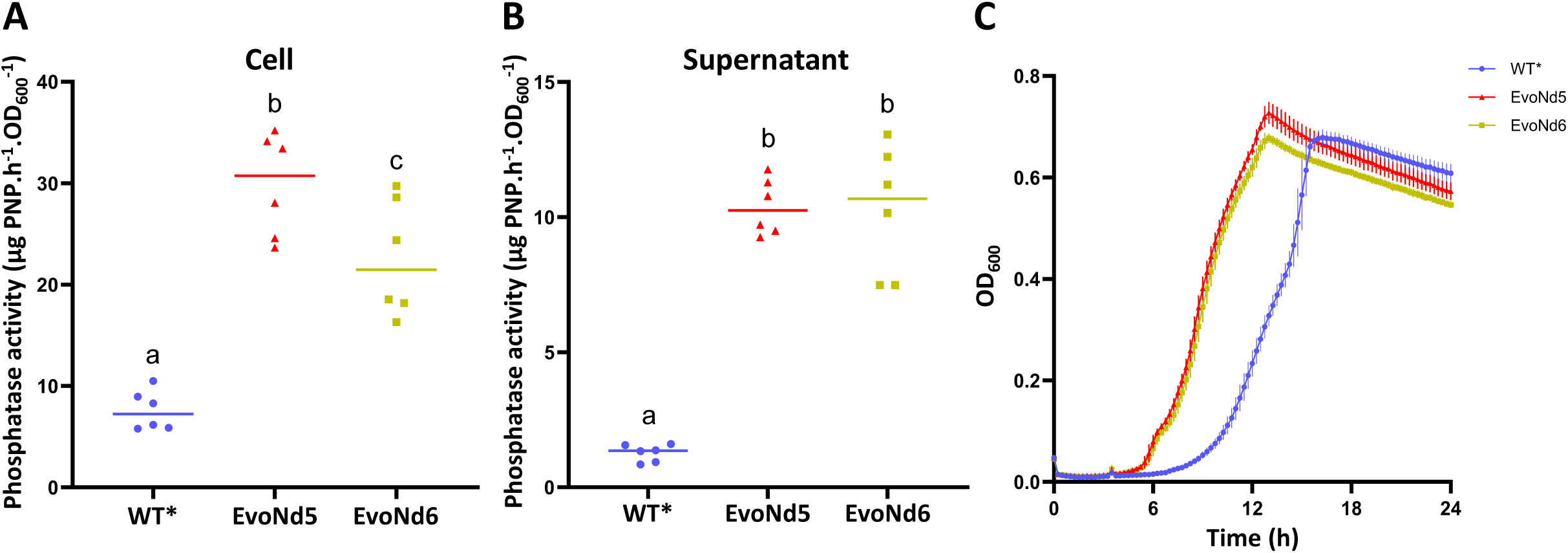
Mutations in *uxpB* increase phosphatase activity. (A-B) Cell-associated (A) and supernatant (B) phosphatase activities of WT* and evolved strains after 24 h growth in MOPS medium containing glycerol 1-phosphate. Activity was measured by pNPP hydrolysis after 1 h and normalized to culture OD_600_. Each point represents one biological replicate (n = 6) ; bars indicate means. * p < 0.05; *** p < 0.0001; ns, not significant. (C) Growth curves of WT* and evolved strains. Data represent mean ± SD from three biological replicates with technical duplicates.

### 2.5. UxpB cellular localization does not affect REE stress resistance

To determine whether either of these two increased activities contributes to improved Nd tolerance in evolved strains, knockout mutants for the Tat-1 system (i.e. Δ*tatC1*) and the T2SS (i.e. Δ*xcpRST*) were generated. The Tat-1 system mediates UxpB translocation from cytoplasm to periplasm, whereas the T2SS is involved in its export from periplasm to the outer membrane (Figure S6A). Almost no differences were observed between the WT* control strain and the two mutants in terms of growth, tolerance to Nd or cell associated-alkaline phosphatase activity (Figure 4A, B, C; Figure S6B, Table S1). A marginal phosphatase activity was detected in the supernatant of the WT* and Δ*tatC1* strains, whereas no activity was detected in the Δ*xcpRST* strain supernatant. (Figure 4D).

**Fig. 4.**
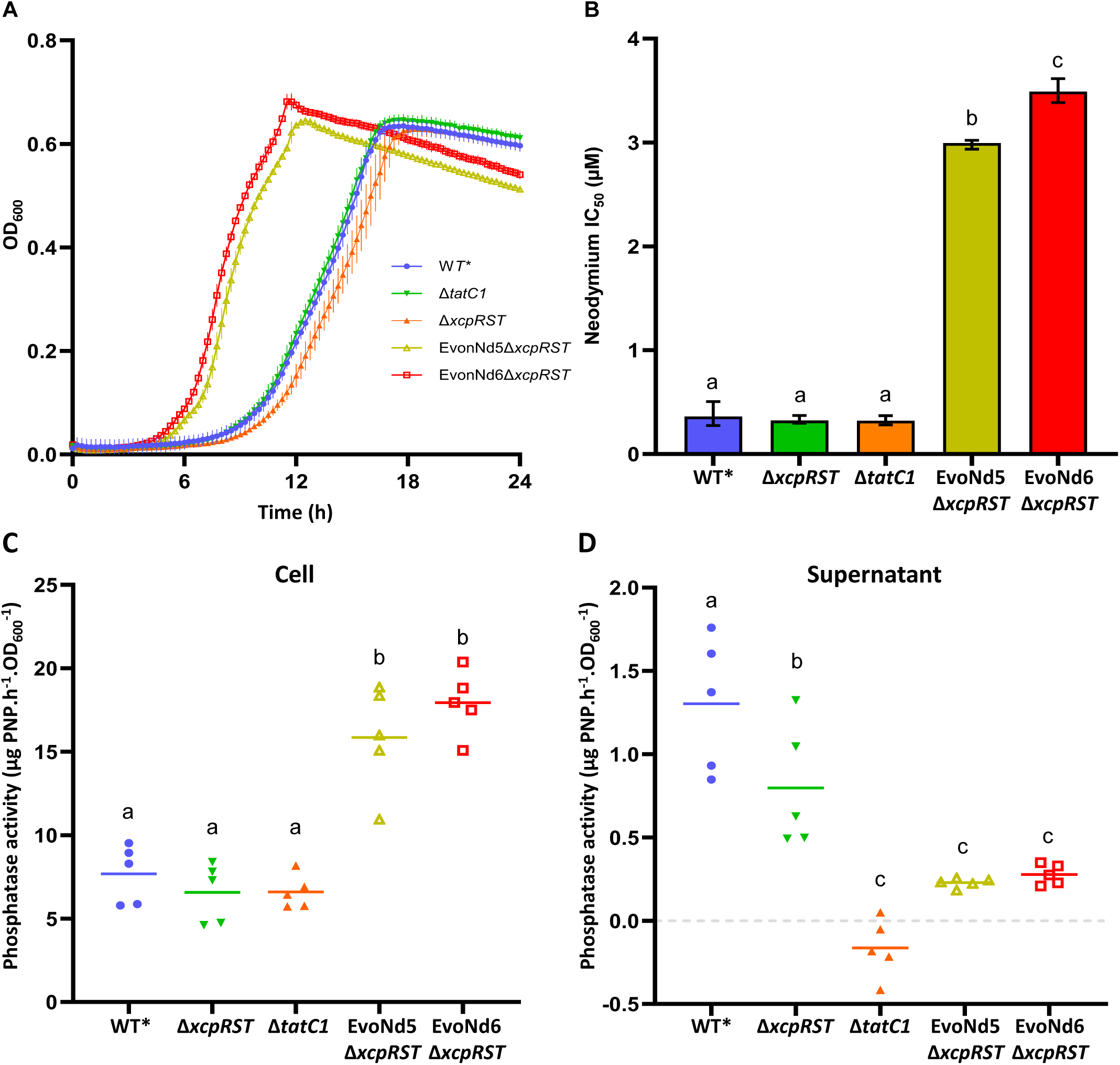
UxpB cellular localization does not influence REE toxicity. (A) Growth curves of WT*, secretion mutants Δ*tatC1* and Δ*xcpRST* and evolution EvoNd5Δ*xcpRST* and EvoNd6Δ*xcpRST* strains in MOPS medium supplemented with glycerol 1-phosphate. Data represent means ± SD from biological triplicates with technical duplicates. (B) Nd IC_50_ values of the indicated strains after 24 h growth. Error bars indicate 95% confidence intervals. Different letters indicate significant differences (p < 0.05). (C-D) Cell-associated (C) and supernatant (D) phosphatase activities after 24 h of growth. Activity was measured by pNPP hydrolysis after 1h and normalized to OD_600_. Each point represents one biological replicate (n = 5); bars indicate means. Different letters indicate significant differences (one-way ANOVA with Tukey’s post hoc test, p < 0.05).

Inactivation of the T2SS in evolved strains led to a 3-hour reduction in lag time compared to the WT* strain, with growth profiles similar to those of their respective parental strains, EvoNd5 and EvoNd6 (Figures 4A and 3D). The IC_50_ values for the EvoNd5Δ*xcpRST* and EvoNd6Δ*xcpRST* strains were 2.99 and 3.49 µM Nd, respectively, indicating greater resistance than the WT* strain and similar resistance to their parental strains with an active T2SS (Figure 2B and Figure 4B). The phosphatase activities associated with EvoNd5Δ*xcpRST* and EvoNd6Δ*xcpRST* cells were approximately 15.8 and 17.9 µg PNP·h⁻¹·OD_600_⁻¹, respectively, representing an increase compared to the WT* strain but a decrease relative to the EvoNd5 strain, and similar activity to the EvoNd6 strain (Figure 3A and Figure 4C). No phosphatase activity was detected in the supernatants of the EvoNd5Δ*xcpRST* and EvoNd6Δ*xcpRST* strains (Figure 4D).

### 2.6. Inorganic phosphate protects *P. putida* against Nd toxicity

To simulate the effect of increased phosphatase activity on the tolerance of *P. putida* WT* to Nd, dose-response growth inhibition assays were performed using combined gradients of Nd and inorganic phosphate (K₂HPO₄, Pi) concentrations in MOPS medium supplemented with glycerol 1-phosphate (Figure 5A). Pi supplementation had no effect on *P. putida* growth in the absence of Nd. In the presence of 0.5 µM Nd, the WT* strain was unable to grow, but addition of 10 µM (or more) Pi fully restored growth. At 1 µM Nd, 20 µM Pi enabled limited growth, while 50 µM Pi restored full growth. When Nd concentrations reached 20 µM or 50 µM, growth was completely inhibited regardless of the Pi concentration (Figure 5A). These results are consistent with *in silico* simulation predicting that Nd-Pi complexation strongly depends on the relative concentrations of both compounds (Figure 5B). In the absence of Pi, Nd remained entirely in its free ionic form. At low Nd levels (0.5 - 1 µM), even small amounts of Pi (≥1 µM) were sufficient to complex almost all available Nd. In contrast, at higher Nd concentrations, complexation efficiency decreased for the same phosphate levels. For example, at 20 µM Nd, only about half of the metal was complexed with 10 µM Pi, while full complexation required higher Pi supplementation.

**Fig. 5.**
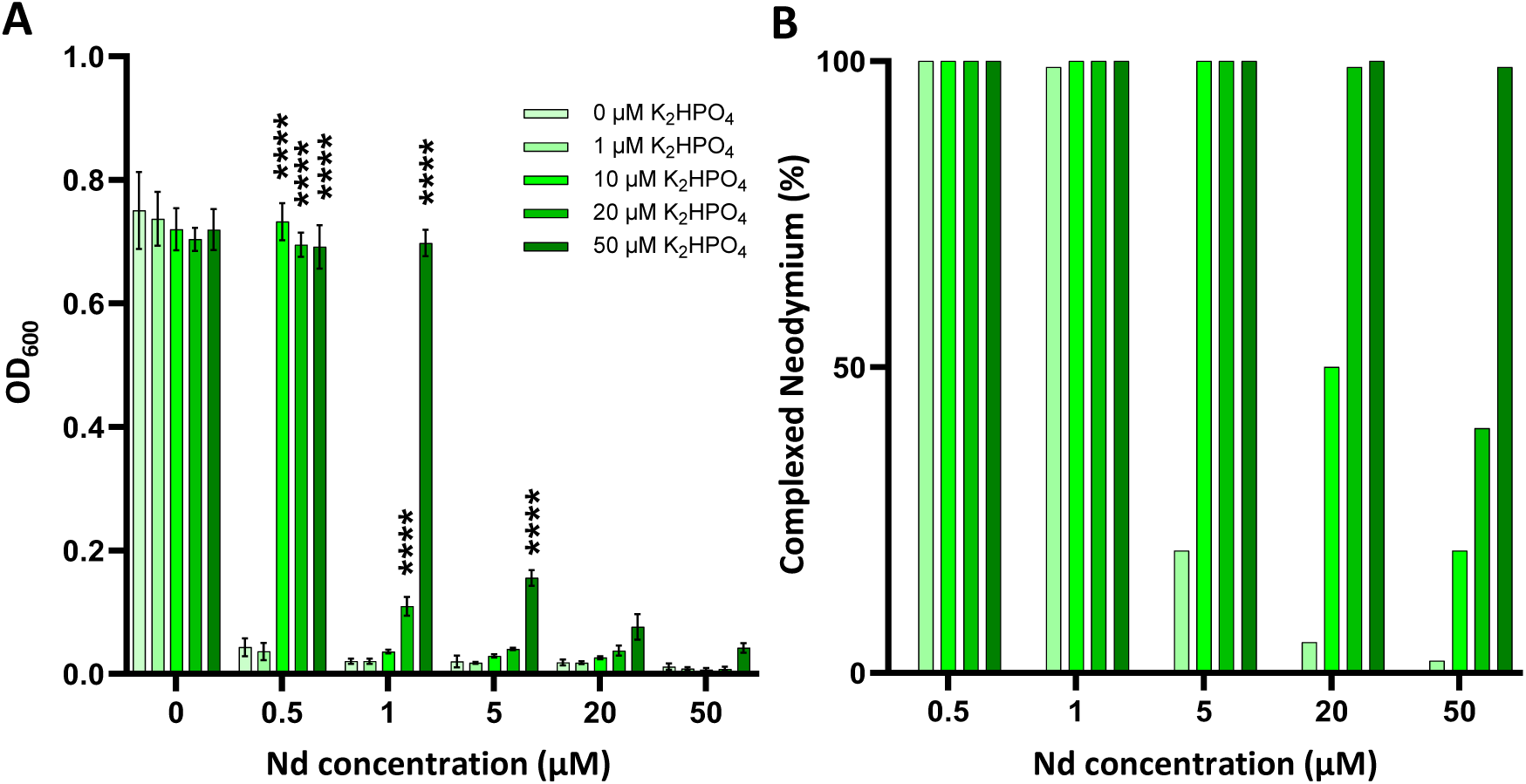
Effect of inorganic phosphate on neodymium toxicity toward *P. putida*. (A) Final growth of *P. putida* WT* after 24 h in MOPS medium containing glycerol 1-phosphate and increasing concentrations of NdCl_3_ (0-50 µM) and/or inorganic phosphate (K_2_HPO_4_, 0-50 µM). OD_600_ values were obtained from five biological replicates (n = 5). Statistical differences relative to the corresponding 0 µM Nd control were assessed using two-way ANOVA followed by Dunnett’s post-hoc test. **** *p* < 0.0001; no annotation indicates no significant difference. (B) Prediction of Nd complexation by phosphate in MOPS medium containing glycerol 1-phosphate, succinate, and increasing K_2_HPO_4_concentrations, calculated using Visual MINTEQ®.

## 3. Discussion

REE are increasingly accumulating in soils and aquatic environments, raising ecotoxicological concerns regarding plants, animals and humans (for reviews, see [8,32]). However, studies on REE toxicity towards bacteria remain scarce. This work examines the toxicity of REE toward *Pseudomonas putida* KT2440, a soil bacterium previously demonstrated to employ LREE as a cofactor of a PQQ-ADH during growth on alcohol substrates. As a first approach to investigate the toxic effects of REE, the strain was cultured on succinate, a preferred carbon sources for *P. putida*. To the best of our knowledge, its metabolism does not depend on these metals. Culture conditions were chosen to ensure high metal bioavailability and accurate assessment of REE effects. Accordingly, *P. putida* was grown in a defined medium buffered with MOPS, generally regarded as a non-complexing agent [33], although weak interactions with REE have been reported [34]. Glycerol 1-phosphate was used instead of inorganic phosphate to avoid metal precipitation while meeting the cell’s nutritional phosphorus requirements. Under these conditions, *P. putida* exhibited high sensitivity to REE with IC_50_ varying around 1 µM. In our experimental setup, these values were several orders of magnitude lower than that of essential metals such as Zn and Ni, and even that of the potent toxicant cadmium (Table 2). This sensitivity is consistent with the low REE concentrations of 1-10 nM supporting REE-dependent growth in the metabolism of alcohols [24]. Efforts to determine whether REE toxicity was altered under these conditions proved inconclusive, as the bacterium was unable to grow on alcohol when glycerol 1-P was supplied as the sole phosphorus source. Although these findings provide important mechanistic insights, IC_50_ values determined in this study are higher than dissolved REE concentrations typically reported in uncontaminated natural environments, where REE are generally found at nanomolar levels. However, such concentrations remain environmentally relevant in anthropogenically impacted settings, including mining areas, industrial effluents, and REE-enriched agricultural soils, where local accumulation can substantially increase REE bioavailability. These concentration ranges therefore, remain valuable for investigating the mechanisms underlying REE toxicity and bacterial adaptive responses.

In addition to global REE sensitivity, the strain exhibited greater sensitivity to LREE than to HREE, as evidence by the strong correlation between IC_50_ values and ionic radii. Identifying such a trend in IC_50_ in relation to physicochemical properties within REE would allow ecotoxicological information derived from a few representatives to be extrapolated to the entire group. However, only a limited number of studies have compared the toxicity of all REE, with contrasting results depending on the model organism studied, the metal form (soluble salts vs particulate) used, and the exposure conditions ([29] and ref. herein). Among those devoted to bacteria, the data from [11] using *E. coli* as a model allows for a relevant comparison with the present work, particularly due to the use of a test medium with a similar composition. In contrast to *P. putida*, a decreasing trend was observed for 24h IC_50_ to *E. coli*, ranging from 18.1 µM for La to 1.1 µM for Sc, with HREE being the most toxic metals. Increased toxicity associated with HREE has also been reported for *Allivibrio fischeri* in bioluminescent based toxicity assays [11,15,35]. These findings suggest that *E. coli* exhibits a generally higher tolerance to REE than *P. putida*, in line with recent observations [36], and may reflect more efficient toxicity mitigation mechanisms. Conversely, the opposite ecotoxicity trends observed across the lanthanides series, as well as for Sc and Y, between the two bacteria warrant further investigation. Our results suggest that the toxicity pattern observed in *P. putida* is unlikely to be explained by a time-dependent decrease in REE exposure concentrations [29]. Instead, we hypothesize that this pattern may arise from differences in the reactivity of individual REE with specific cellular targets and metabolites [37]. This differential reactivity may be related to intrinsic physicochemical differences among REE. According to the Hard–Soft Acid–Base (HSAB) theory, LREE are more polarizable soft acids, whereas HREE behave as hard acids, a difference that influences how they interact with biological ligands: LREE tend to form stronger inner-sphere complexes with soft, polarizable donor atoms (e.g., carboxylate, phosphate), while HREE prefer harder, less polarizable donors. The cellular responses to REE may thus vary depending on the specific metal-ligand interactions dictated by the biochemical environment and dominant functional groups of each cell. In this context, integrative omics approaches provide a powerful means to identify REE-altered metabolic pathways and cellular processes and to elucidate their underlying mechanisms of action.

As a first step in studying theses mechanisms, selection of *P. putida* under continuous exposure to gradually increasing REE concentrations yielded two variants showing 7- and 10-fold enhanced tolerance to REE. We found that the causative mutations are substitutions within the alkaline phosphatase encoding gene *uxpB*, and more specifically in its signal sequence region. UxpB is a lipoprotein whose secretion and modification pathway in *P. putida* has been documented [30]. The cell employs two Tat secretion systems (Tat-1 and Tat-2) for UxpB export under low and high phosphate conditions, respectively. After translocation to the periplasm, UxpB undergoes lipid modification through the canonical Lgt– Lsp–Lnt pathway: Lgt attaches a diacylglycerol, Lsp removes the signal peptide, and Lnt transfers an additional acyl group to the N-terminus [38]. The mature, acylated UxpB is then secreted via a type II secretion system (T2SS), a multiprotein complex (XcpA and XcpP– XcpZ) spanning the cell envelope that drives substrate export through a pseudopilus-mediated mechanism [39,40]. Once secreted, UxpB is anchored in the outer membrane via its acyl chain, but can be released into the extracellular medium depending on the growth conditions [30]. Our deletion and overexpression experiments revealed a correlation between *uxpB* expression, cell associated phosphatase activity and tolerance to Nd. One possible explanation is that the enhanced release of phosphate resulting from glycerol 1-phosphate hydrolysis by UxpB promotes the formation of Nd-Pi complexes, which may reduce Nd bioavailability and consequently its toxicity

Interestingly, phosphatase activity was detected in the culture supernatant of the WT* strain, but not when *uxpB* was expressed *in trans* to complement the Δ*uxpB* mutant. This observation may reflect saturation or imbalance in the Tat/T2SS secretion machinery or altered UxpB targeting caused by phosphatase overexpression. Nevertheless, this imbalance did not prevent the improvement in Nd tolerance. The study of the evolved strains EvoNd5 and EvoNd6 indicated that their increased tolerance to Nd is linked to elevated UxpB activity. Both strains showed a higher phosphatase activity and a reduction of the lag time compared to WT*, consistent with improved phosphate acquisition. Although increased phosphatase activity was also detected in culture supernatants, extracellular activity was not required for REE tolerance, as mutants lacking the T2SS secretion and modification system displayed unchanged Nd tolerance. Similarly, inactivation of the Tat1 system and T2SS in the parental WT* strain did not affect basal Nd tolerance. Together, these results indicate that periplasmic UxpB activity is sufficient and likely represents the dominant mechanism contributing to REE tolerance. Consistently, exogenous phosphate addition increased Nd tolerance in a dose-dependent manner, supporting a reduction of REE bioavailability through phosphate complexation. Interestingly, the evolved strains also displayed a modest but significant increased tolerance to Cd and Zn, consistent with a similar phosphate mediated reduction in metal bioavailability. In this case, the more limited gain in tolerance could be due to the lower tendency of these metals to precipitate in the presence of phosphate compared to REE (calculated solubilities of Zn (PO) and Cd (PO) of 1.33×10 mol·L ¹ and 1.03×10 mol·L ¹ respectively, versus 1.95×10 ¹³ mol·L ¹ for NdPO [41,42]. Although mechanistic analyses were mainly performed with Nd, similar interactions with phosphate are likely to occur for other REE because of their shared chemical properties. However, differences in coordination chemistry and phosphate solubility across the lanthanide series may influence the extent of these effects. Since REE immobilization was not directly demonstrated here, further work will be required to assess the generality of this mechanism.

Previous studies linked bacterial tolerance to toxic metals to phosphatase-mediated release of inorganic phosphate and subsequent precipitation of metal-phosphate minerals. For instance, *Caulobacter crescentus* lacking the periplasmic phosphatase PhoY showed impaired U(VI) precipitation and markedly reduced survival, demonstrating a causal relationship between biomineralization and metal tolerance [43]. Similar correlations have been reported in *Achromobacter xylosoxidans*, where Pb exposure induces phosphatase activity and leads to the formation of protective pyromorphite precipitates [44], and in *Citrobacter* mutants overproducing phosphatase, which accumulate metals more efficiently and exhibit enhanced tolerance [45]. Beyond its links to metal tolerance, microorganism-induced phosphate precipitation is also gaining importance as an effective tool for metal bioremediation. [46]. To our knowledge, the present study provides for the first time compelling evidence that REE could be bio-immobilized through the phosphatase activity of *P. putida*, highlighting its potential to be used for REE bioremediation and biorecovery [47].

## 4. Conclusion

Overall, this study reveals a marked sensitivity of *Pseudomonas putida* KT2440 to soluble REE and highlights the importance of species-specific determinants in bacterial responses to these metals. Through experimental evolution and genetic analysis, we identify alkaline phosphatase UxpB as a key driver of REE tolerance, with mutations enhancing its activity leading to increased phosphatase-dependent release of inorganic phosphate and subsequently, probable precipitation of REE-phosphate complexes. These findings strongly suggest that *P. putida* can mitigate REE toxicity via biomineralization, consistent with phosphatase-mediated metal immobilization pathways previously described in other bacteria. Beyond improving our understanding of REE-microbe interactions, this work also demonstrates that enhanced phosphatase activity can substantially reduce REE bioavailability and toxicity, potentially providing a mechanistic basis for developing *P. putida* as a chassis for REE bioremediation or biorecovery from secondary resources. In line with the growing recognition of microbial induced phosphate precipitation as an effective tool for metal remediation [46] and the increasing interest in microbial strategies for REE recovery [47], our findings position *P. putida* as a promising candidate for future applications targeting REE-contaminated environments and waste streams.

## 5. Methods

### 5.1. Bacterial strains, plasmids and culture conditions

A list of strains and plasmids used in this study is provided in Table S2. For phenotypical assays, *Pseudomonas putida* KT2440 and derivatives were cultured in a modified MOPS medium [48]: MOPS 40 mM, NH_4_Cl 9.52 mM, MgCl_2_ 1 mM, K_2_SO_4_ 276 µM, FeSO_4_ 10 µM, CaCl_2_ 50 µM, NaCl 50 mM, Tricine 4 mM, sodium succinate dibasic hexahydrate 25 mM, rac-glycerol 1-phosphate sodium 1 mM (Sigma). If necessary, gentamycin 30 µg.mL^-1^ and L-arabinose (Sigma) 0.1% w/v or 0.0125% was added. If not stated otherwise, overnight cultures were performed in the same medium as the culture conditions. Growth was performed at 28°C, 180 rpm with an initial OD_600_ of 0.05. All REE were purchased from Sigma in their chloride form. REE solutions were prepared in double-distilled water and were sterilized by microfiltration.

For plasmids and mutants constructions, *P. putida* and *Escherichia coli* were grown in lysogenic broth (LB, Sigma) or M9 medium [49]. If necessary, kanamycin 50 µg/ml, gentamicin 30 µg/ml, or 5-fluorouracil 20 µg/ml was added. Agar at 15 g.L^-1^ was added for solid media.

### 5.2. Dose-response growth inhibition assays

Strains were cultured in a 96 well-plate in the presence of a concentration gradient of each metal, ranging from 9.7 nM to 10 µM, prepared by serial dilution. Final OD_600_ were measured after 24h of incubation and results were normalized using the absorbance of a REE-free culture as the 100% value. Half maximal growth inhibition (IC_50_) values were calculated using four parameters non-linear regression (Prism version 10.4.2). If data were not sufficient to calculate 95% confidence interval, supplementary growth inhibition assays were conducted using REE concentration above the hypothetical IC_50_ until confidence interval was obtained. Statistical differences between dose-response curves were assessed using extra sum-of-square F test (Prism version 10.4.2).

### 5.3. REE analyses

REE quantification in samples was performed by inductively coupled plasma mass spectrometry (Thermo iCap TQ ICP-MS,Thermo Fisher). ICP-MS instrument was tuned daily for oxide and doubly charged ions and mass calibration accuracy was checked. SQ-KED (Single Quadrupole-Kinetic Energy Discrimination) or TQ-O2 mass-shift modes (Triple Quadripole Oxygen Reaction) were used to limit potential polyatomic or isobaric interferences. External calibration was performed using synthetic multi-elemental solution (AccuStandard) acidified with 2% HNO_3_ (v/v) (NORMATOM, VWR). The accuracy of the calibration was checked with certified reference surface water (SPS-SW1, Spectrapure Standards). The instrumental drift was monitored by running SPS-SW1 every 10–20 samples and by adding Rh and Ir to the samples as internal standards. The measurements were performed three times and averaged for each solution.

Calculation of REE speciation in the culture medium was carried out with Visual MINTEQ 3.1 software using the default database. In addition, relevant stability constants of REE ligands complexes and solubility products for REE phosphates mineral phases were retrieved from those selected by [50]. The input concentration for each component was calculated from MOPS medium composition.

### 5.4. Isolation of REE stress-tolerant variants

The *P. putida* KT2440 WT* strain was grown in 96-well plate in the presence of various NdCl_3_ concentrations (from 250 nM to 2 µM) for 24h. Wells showing bacterial growth at the highest Nd concentrations were used to restart a growth cycle under the same conditions. This procedure was repeated until a stable Nd-resistant phenotype was obtained. Dilution of these cultures were then plated on non-selective LB plates and individuals clones were further streaked on LB plate before their resistance phenotype was assessed by IC_50_ determination. Evolved strains with the higher resistance toward Nd were selected and the causative mutations were identified by Oxford Nanopore sequencing of their genome (PlasmidSaurus, Louisville, KY, USA) and variant calling. Sequences were aligned to the *P. putida* KT2440 reference genome (accession number: AE015451.2) using BWA MEM, and single nucleotide polymorphism (SNP) calling was performed using FreeBayes and BCFtools, available via the BV-BRC website [51].

### 5.5. General molecular biology procedures

High fidelity polymerase chain reaction (PCR) was performed using Fidelio Hot Start DNA polymerase (Ozyme). Control PCR was performed using Phire Hot Start PCR master mix (Thermo Scientific). Primer used is this study are listed in Supplementary Table 3. Restriction enzymes were used according to manufacturer’s instructions (Thermo Scientific). Genomic DNA extraction and purification were performed using the Wizard Genomic DNA Purification Kit (Promega). Nucleospin Plasmid kit and Nucleospin Gel and PCR Clean-up kit (Macherey Nagel) were used for plasmid extraction and PCR products purification, respectively. Gibson assembly was performed using the NEBuilder Hifi DNA Assembly Master Mix (New England Biolabs). All kits were used according to manufacturers’ instructions.

### 5.6. Construction of plasmids

For construction of deletion plasmids pJOE-uxpB and pJOE-xcpRST the regions around 800-1000 bp upstream and downstream of genes *uxpB* and *xcpRST* were amplified from KT2440 genomic DNA by PCR using primer pairs Up_*uxpB*_fwd/Up_*uxpB*_rev and Down_*uxpB*_fwd/Down_*uxpB*_rev, Up_*xcpR*_fwd/Up_*xcpR*_rev and Down_*xcpT*_fwd/Down_ *xcpT*_rev, respectively. The up-and dowstream amplicons were then assembled via Gibson assembly together with BamH1 digested pJOE6261.2 and transformed into *E. coli* DH5_α_. The previously constructed plasmid pJOE-tatC1 [27] was used for deletion of the *tatC1* gene.

To construct the expression plasmid pJN105:*uxpB*, the *uxpB*_fwd/*uxpB*_rev primer pair was used to amplify the wild-type *uxpB* gene. The pJN105 vector was digested by XbaI and joined with the amplicon by Gibson assembly so that the expression of *uxpB* is under the control of an L-arabinose inducible promoter [52].

### 5.7. Construction of deletion strains

For in-frame deletion of chromosomal genes, we employed the previously established method of [53]. Briefly, following electrotransformation [54] with deletion plasmids, clones resistant to kanamycin (Kan) and sensitive to 5-fluorouracil (5-FU) were selected. One clone was subsequently incubated in liquid LB medium for 24 h at 30° C with shaking at 180 rpm. Afterward, selection for 5-FU resistant and Kan sensitive clones was performed on M9 minimal medium plates, and clones carrying the desired gene deletion were identified by colony PCR.

### 5.8. Phosphatase activity measurement

Phosphatase activity was measured by monitoring degradation of *para*-nitrophenyl phosphate (*p*NPP) into *para*-nitrophenol (PNP). Measurements were performed either with cells after one wash in sterile growth medium or with supernatant collected after centrifugation (7000 g x 2 min). 480 µL of cells or supernatant was supplemented with 20 µL of 100 mM *p*NPP solution (Sigma; final concentration 4 mM) and the mixture was incubated 1h at room temperature. Next, 50 µL of a 1 M NaOH solution (final concentration 0.1 mM) was added, followed by 10 minutes of incubation at room temperature. After centrifugation (8000 x g, 2 minutes), the supernatant was collected and the 405 nm optical density (OD) was measured. The amount of PNP produced was determined using a standard curve ranging from 0 to 100 µg.mL^-1^ of purified PNP (Sigma). The results are expressed in µg of PNP synthesized per hour and are normalized with the OD_600_ of each culture.

## CrediT authorship contribution statement

**Charly A. Dupont:** Conceptualization, Data curation, Investigation, Methodology, Writing – original draft, Writing – review and editing. **Theophile Franzino**. Investigation, Writing – review and editing. **Lola Perey**. Investigation, Writing – review and editing. **Maximilien Beuret.** Investigation, Writing – review and editing**. Nolhan Berceaux**. Investigation, Writing – review and editing. **Patrick Billard**. Conceptualization, Data curation, Funding acquisition, Investigation, Methodology, Supervision, Validation, Writing – original draft, Writing – review and editing.

## Funding

This work was supported by the Agence Nationale pour la Recherche under grant ANR-23-CE44-0016 and under the France 2030 program, reference ANR-22-PERE-0003 for C.A. Dupont.

## Declaration of Competing Interest

The authors declare that they have no known competing financial interests or personal relationships that could have appeared to influence the work reported in this paper.

## Acknowledgements

We thank Hélène Le Cordier (LIEC) for the technical help provided. We also thank the pôle de compétences en chimie analytique, ANATELo (LIEC) and Jose Paulo Pinheiro for his assistance in using Visual MINTEQ.

## Notes

### Competing Interest Statement

The authors have declared no competing interest.

### Summary of Updates

Missing Table 1 and Table 2 were added.

