## Supplementary Files for "Phosphatase-mediated mitigation of rare earth element toxicity to *Pseudomonas putida*"

**New insights into the toxicity of rare earth elements in *Pseudomonas putida* KT2440.**

**Table of contents**

**Table S1:** IC_50_ parameters of Nd toward *Pseudomonas putida* mutants.

**Table S2:** Strains and plasmids used in this study.

**Table S3:** Oligonucleotides used in this study

**Figure S1:** Dose-response curves of *P. putida* WT* exposed to rare earth elements.

**Figure S2:** Dosage of selected rare earth elements in the experimental conditions.

**Figure S3:** Dose-response curves of WT* and evolved strains exposed to zinc, nickel and cadmium.

**Figure S4:** Dose-response curves of evolved strains exposed to rare earth elements.

**Figure S5:** Dose-response curves of *P. putida* WT* + EV and Δ*uxpB* + *uxpB* strains exposure to Nd.

**Figure S6:** Dose-response curves of *P. putida* Δ*xcpRST*, Δ*tatC1*, EvoNd5Δ*xcpRST*, EvoNd6Δ*xcpRTS* strains to neodymium exposure.

**Table S1:** IC_50_ parameters of Nd toward *Pseudomonas putida* mutants.

Numerical values of the IC_50_ and the associated 95% confidence interval (CI) for figures 1B, 2B and 4B.

| **Strain** | **Nd IC_50_ (µM)** | **95% CI** |
| --- | --- | --- |
| WT* | 0.360 | 0.274 to 0.504 |
| EvoNd5 | 3.863 | 3.829 to 3.917 |
| EvoNd6 | 2.792 | 2.756 to 2.883 |
| WT* + EV | 0.683 | 0.489 to 0.946 |
| Δ*uxpB* + *uxpB* - 0% L-ara | 0.269 | 0.243 to 0.296 |
| Δ*uxpB* + *uxpB* - 0.0125% L-ara | 0.960 | 0.880 to 1.054 |
| Δ*uxpB* + *uxpB* - 0.1%  L-ara | 2.697 | 2.609 to 2.792 |
| Δ*xcpRST* | 0.323 | 0.298 to 0.371 |
| Δ*tatC1* | 0.321 | 0.281 to 0.370 |
| EvoNd5Δ*xcpRST* | 2.996 | 2.936 to 3.020 |
| EvoNd6Δ*xcpRST* | 3.490 | 3.384 to 3.615 |

**Table S2:** Strains and plasmids used in this study.

| **Strains** | **Relevant Characteristics** | **Reference/Source** |
| --- | --- | --- |
| *Pseudomonas putida* |  |  |
| KT2440 | Wildtype strain of *Pseudomonas putida* (ATCC 47054). | ATCC 47054 |
| Δ*upp* (WT*) | KT2440 with a *in frame* deletion of *upp.* Parental strain for gene deletions. | Graf and Altenbuchner, 2011 |
| EvoNd5 | Neodymium resistant WT* strain. Ponctual mutation leading to the G5S substituion in UxpB. | This study |
| EvoNd6 | Neodymium resistant WT* strain. Ponctual mutation leading to the G47S substituion in UxpB. | This study |
| Δ*uxpB* | ∆*upp* with a markerless mutation of *uxpB*. | This study |
| Δ*tatC*1 | ∆*upp* with a markerless mutation of *tatC1*. | This study |
| Δ*xcpRST* | ∆*upp* with a markerless mutation of *xcpR, xcpS* and *xcpT.* | This study |
| Δ*upp*  + pJN105 | Δ*upp* strain with pJN105 empty vector, GmR. | This study |
| Δ*uxpB* + pJN105 | Δ*uxpB* strain  with pJN105 empty vector, GmR. | This study |
| Δ*uxpB* + pJN105:*uxpB* | Δ*uxpB* strain  with pJN105 carrying wild-type *uxpB* gene, GmR. | This study |
| *Escherichia coli* |  |  |
| DH5α | *fhuA2 lac(del)U169 phoA glnV44 Φ80' lacZ(del)M15 gyrA96 recA1 relA1 endA1 thi-1 hsdR17* | ThermoFischer Scientific |
| **Plasmids** |  |  |
| pJOE6261.2 | Suicide vector for gene deletions. Derivative vector of the pIC20HE vector carrying the *upp* gene, KmR, AmpR. | Graf and Altenbuchner, 2011 |
| pJOE6261.2:Δ*uxpB* | pJOE6261.2 vector carrying the sequence used for *uxpB* gene *in-frame* deletion | This study |
| pJOE6261.2:Δ*tatC1* | pJOE6261.2 vector carrying the sequence used for *tatC1* gene *in-frame* deletion | This study |
| pJOE6261.2:Δ*xcpRST* | pJOE6261.2 vector carrying the sequence used for *xcpRST* genee *in-frame* deletion | This study |
| pJN105 | Arabinose-inducible cloning vector, derivative of pBBR1-MCS5, GmR. | Newman et al., 1999 |
| pJN105:*uxpB* | pJN105 plasmid carriyng the *uxpB* gene under control of an L-arabinose inducible promoter. GmR. | This study |

**Table S3:** Oligonucleotides used in this study

| **Mutagenesis primers** | **Sequences (5’ to 3’)** |
| --- | --- |
| *uxpB deletion* |  |
| Up_*uxpB*_fwd | cgatggccgctttggtcccggcgttgtgcgtctgccag |
| Up_*uxpB*_rev | tgaccggcacccgcgactttaaggcgcaag |
| Down_*uxpB*_fwd | aaagtcgcgggtgccggtcagccaacttttg |
| Down_*uxpB*_rev | cctgcaggtcgactctagagagcgccagttgcgttacttac |
| *tatC1 deletion* |  |
| Up_*tatC1*_fwd | cgatggccgctttggtcccgcccatccgtgcatgcctc |
| Up_*tatC1*_rev | cgaaagggccgaagcattttccttgggcag |
| Down_*tatC1*_fwd | aaaatgcttcggccctttcgcgggcgtg |
| Down_*tatC1*_rev | cctgcaggtcgactctagagggccatgccgagttcgcc |
| *xcpRST deletion* |  |
| Up_*xcpRST*_fwd | cgatggccgctttggtcccggcagtaacggggtgaatatc |
| Up_*xcpRST*_rev | aaggggtgccatagggcagcatcaccag |
| Down_*xcpRST*_fwd | gctgccctatggcaccccttatcagtacc |
| Down_*xcpRST*_rev | cctgcaggtcgactctagagatcagttcgatcagggtcag |
| **Overexpression primers** | **Sequences (5’ to 3’)** |
| *uxpB overexpression* |  |
| uxpB_fwd | cccgtttttttgggctagcgaggaggcaccctgatgagtcgagata |
| uxpB_rev | tggatcccccgggctgcagggtacagccagcaaaagttg |

**Supplementary Figures**

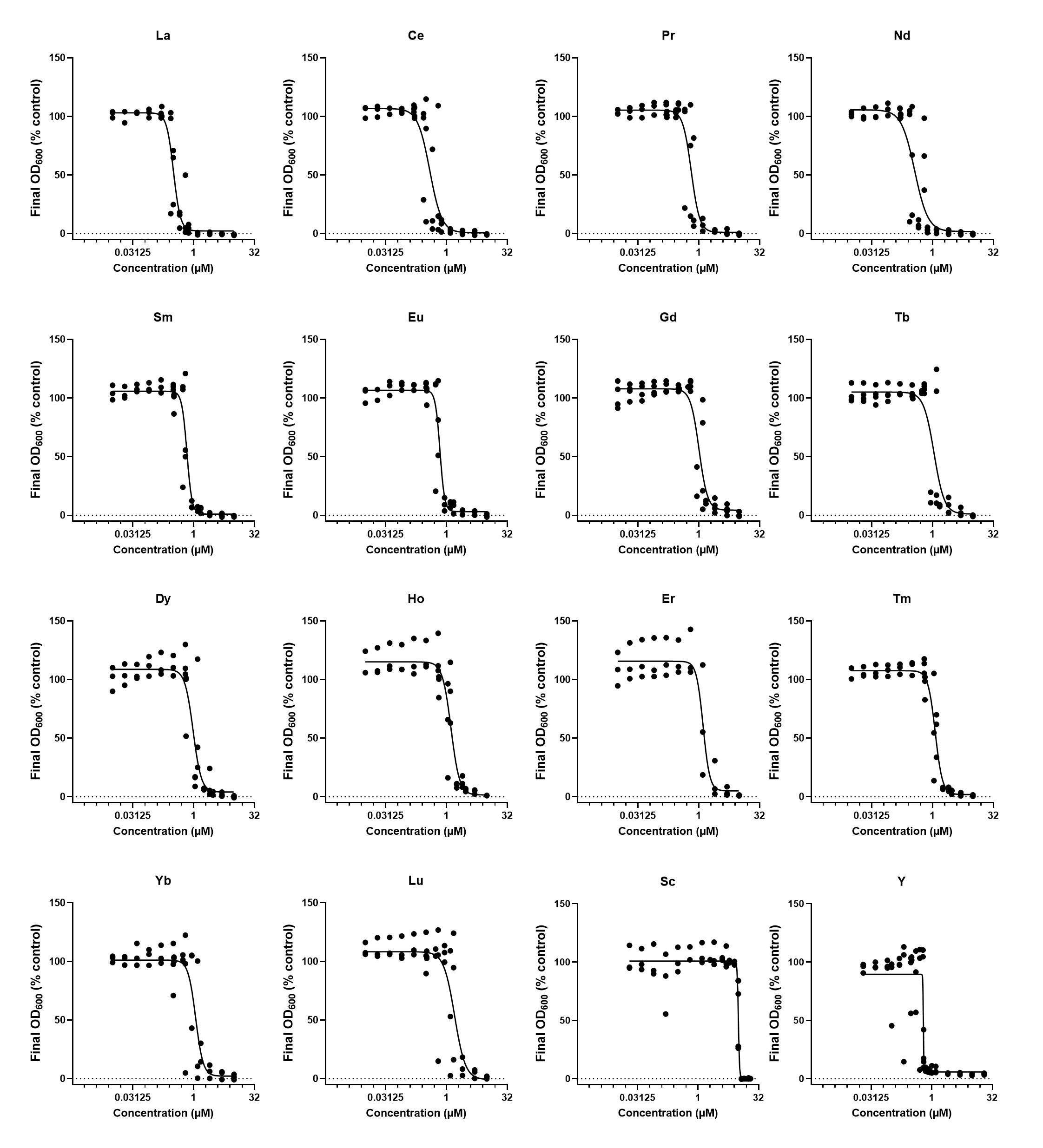

**Figure S1:** Dose-response curves of *P. putida* WT* exposed to rare earth elements. Assays were conducted in MOPS medium with glycerol 1-phosphate as a phosphorus source. Each point represents a biological replicate. Lines represents the best-fit dose-response curve obtained from non-linear regression. Data were obtained after 24h of incubation at 28°c and 180 rpm in biological triplicates (n=3) with technical duplicates.

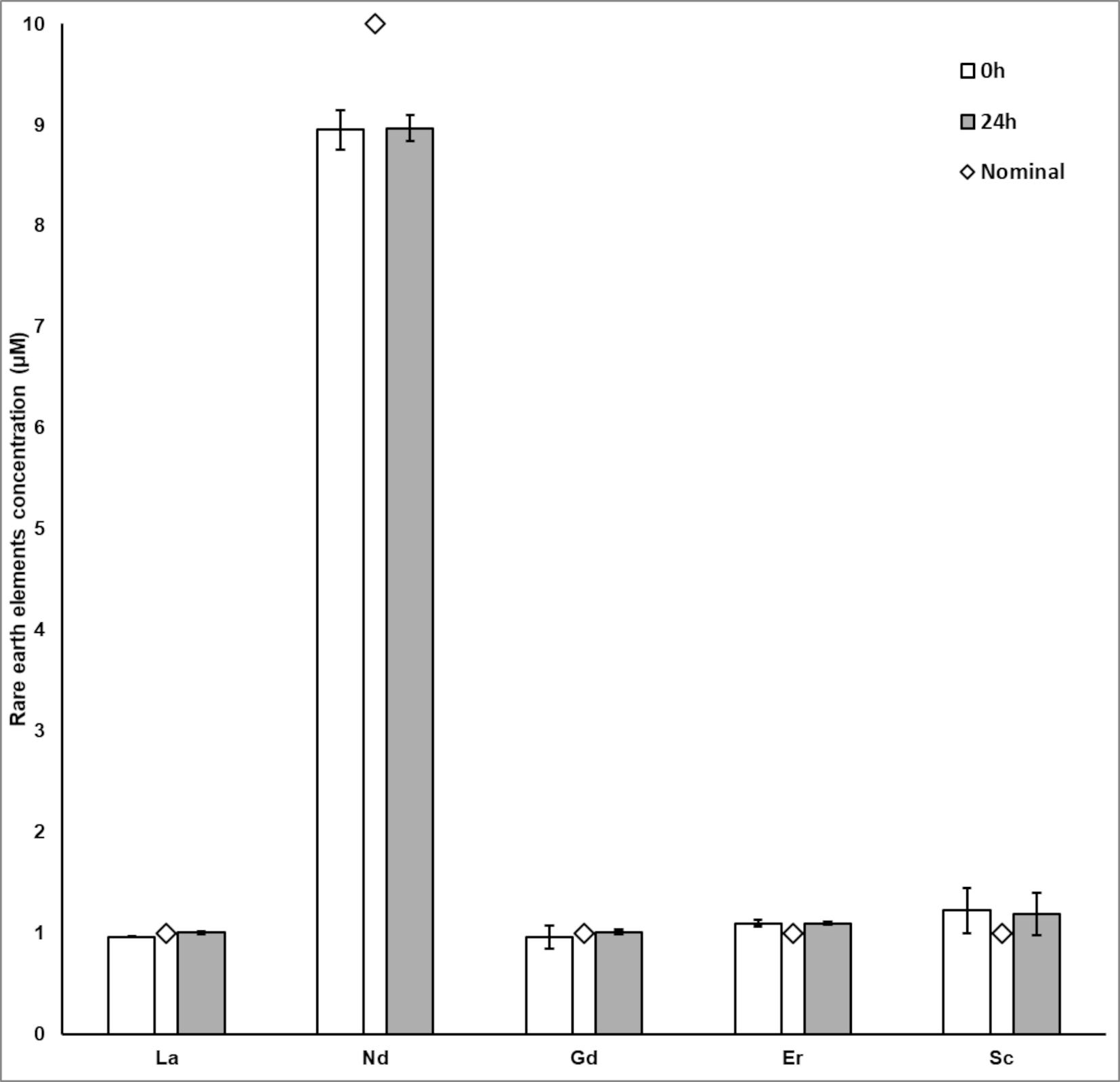

**Figure S2:** Quantification of selected rare earth elements under experimental conditions.

The concentrations of lanthanum (La), neodymium (Nd), gadolinium (Gd), erbium (ER) and scandium (Sc) were determined immediately after addition (0 h, white box) and after 24 h of incubation (grey box). The diamonds represent the nominal concentration.

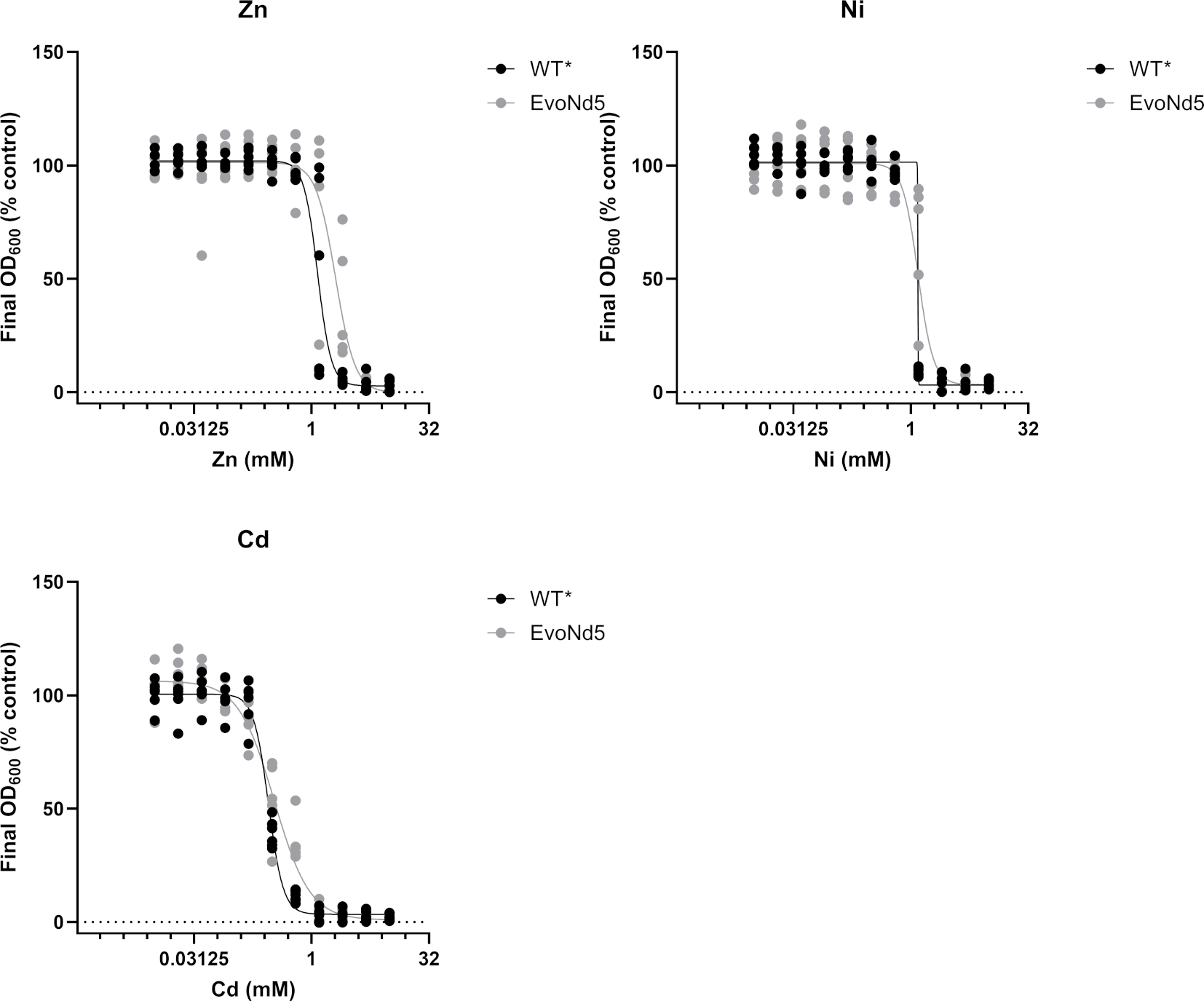

**Figure S3:** Dose-response curves of WT* and evolved strains exposed to zinc, nickel and cadmium.

Assays were conducted in MOPS medium with glycerol 1-phosphate as a phosphorus source. Each point represents a biological replicate. Lines represents the best-fit dose-response curve obtained from non-linear regression. Data were obtained after 24h of incubation at 28°c and 180 rpm in biological triplicates (n=3) with technical duplicates.

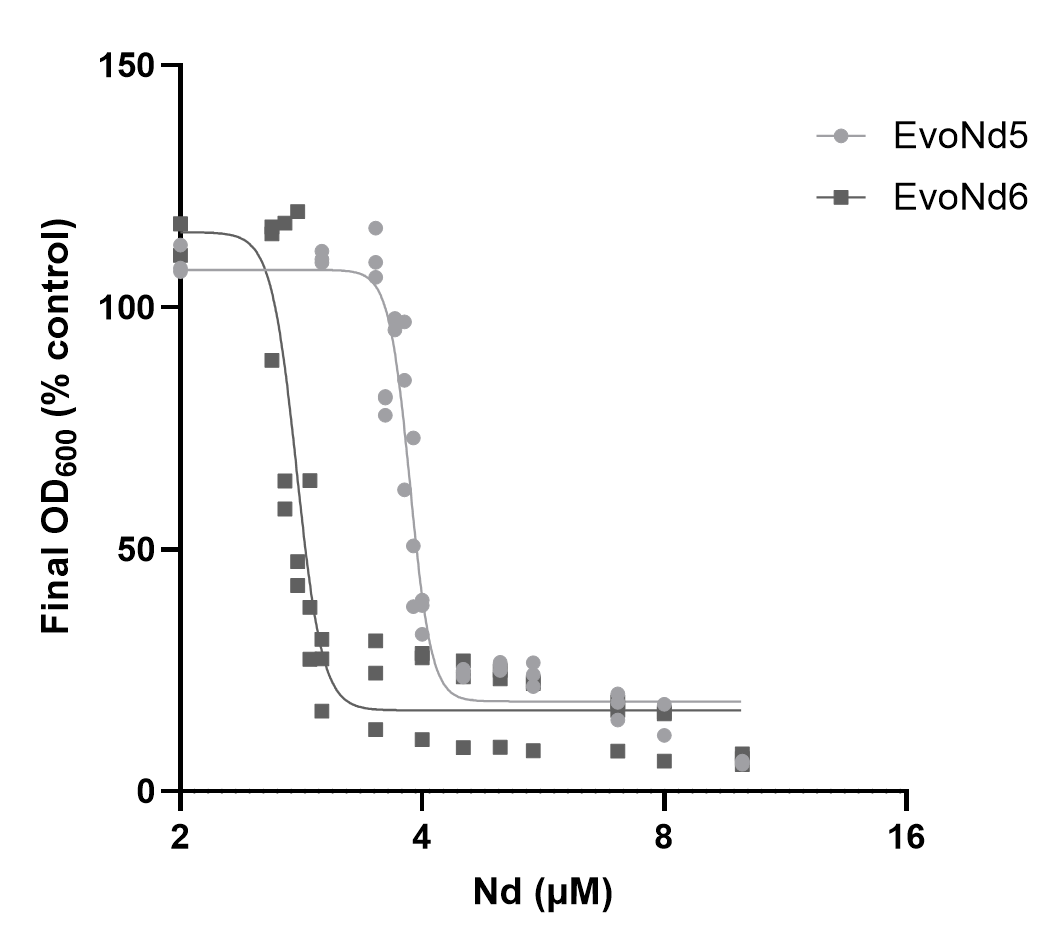

**Figure S4:** Dose-response curves of evolved strains exposed to rare earth elements. Mutants were grown in MOPS medium with various concentrations (0 to 10 µM) of neodymium and their OD_600_ was monitored after 24 h of growth at 28°C and 180 rpm. Each point represents a biological replicate. Lines represents the best-fit dose-response curve obtained from non-linear regression. Data were obtained after 24h of incubation in biological triplicates (n=3) with technical duplicates.

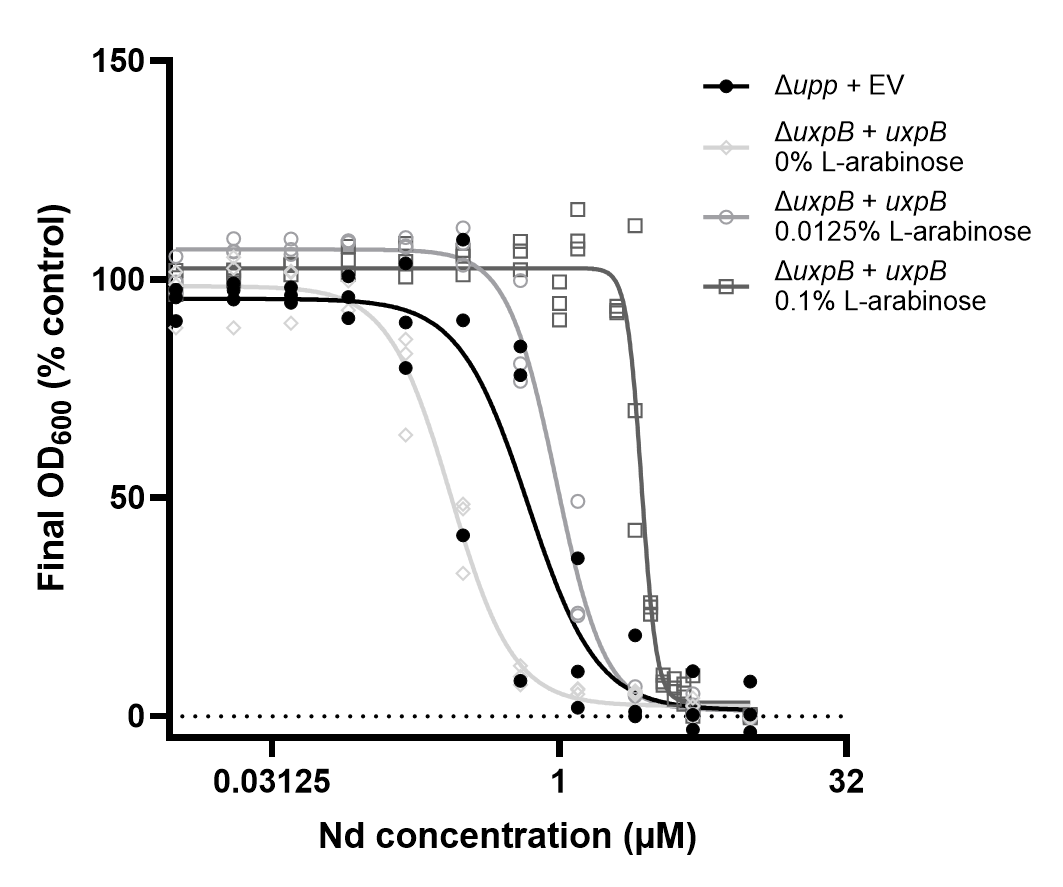

**Figure S5:** Dose-response curves of *P. putida* WT*+ EV and Δ*uxpB* + *uxpB* strains exposure to Nd. Strains were grown in MOPS medium with 0 to 10µM Nd. The Δ*uxpB*+*uxpB* strain was grown with 0%, 0.0125% or 0.1% L-arabinose. OD_600_ were monitored after 24 h of growth at 28°C and 180 rpm. Lines represents the best-fit dose-response curve obtained from non-linear regression. Data were obtained after 24h of incubation in biological triplicates (n=3) with technical duplicates. Black line, black filled circles: WT*+EV. Gray lines, Δ*uxpB*+*uxpB*. Empty diamonds, 0% L-arabinose. Empty circles, 0.0125% L-arabinose. Empty squares, 0.1% L-arabinose.

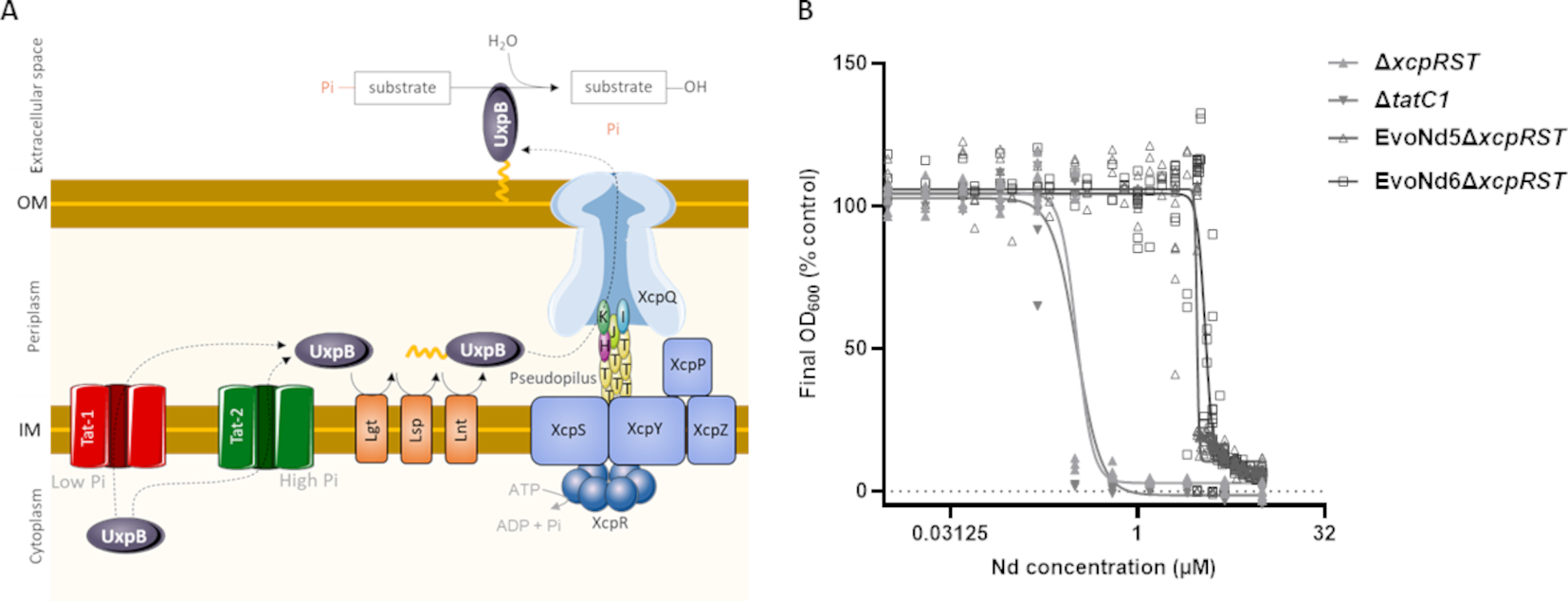

**Figure S6:** Dose-response curves of *P. Putidaa* Δ*xcpRST*, Δ*tatC1*, EvoNd5Δ*xcpRST*, EvoNd6Δ*xcpRTS* strains to neodymium exposure. **A:** Overview of UxpB export, post-translational modifications and function in *P. putida* KT2440. UxpB prelipoprotein is translocated from cytoplasm to periplasm through the Twin-Arginine-Translocation 1 system (Tat) under low phosphate concentration or the Tat 2 system under high phosphate concentration. Once in periplasm, UxpB is acylated by the prelipoprotein diacylglyceryl transferase (Lgt), the lipoprotein signal peptidase II (Lsp) and the lipoprotein *N*N-acyl transferase (Lnt) enzymes. When acylated, the UxpB lipoprotein is translocated to the extracellular space through the Type II Secretion System (T2SS, Xcp). Once secreted, UxpB is anchored to the outer membrane (OM) by its acyl part and can hydrolyse organophosphorus substrates, releasing bioavailable phosphate. Pi: phosphate. IM: inner membrane. **B:** Assays were conducted in MOPS medium with glycerol 1-phosphate as a phosphorus source. Strains were grown with 0 to 10µM Nd. OD_600_ were monitored after 24 h of growth at 28°C and 180 rpm. Lines represents the best-fit dose-response curve obtained from non-linear regression. Each point Data obtained after 24h of incubation in biological triplicate (n=3) with technical duplicate. Filed triangle, ∆*xcpRST*. Filed inverted triangle, ∆*tatC1*. Empty triangle, EvoNd5∆*xcpRST*. Empty square, EvoNd6∆*xcpRST*.
